# The Spatial Specificity and Recovery from Visual Adaptation in Causality Perception

**DOI:** 10.64898/2026.04.07.716922

**Authors:** Laura van Zantwijk, Martin Rolfs, Sven Ohl

**Affiliations:** Department of Psychology Humboldt-Universität zu Berlin, Rudower Chaussee 18, 12489 Berlin, Germany

**Keywords:** perception of causality, visual adaptation, recovery, spatial specificity

## Abstract

When one object approaches another object which, upon touching, moves in the same direction, humans report a vivid impression of one launching the other. Visual adaptation can alter this perception of causality: observers less often report seeing a launch after viewing a stream of launch events. In three experiments, we further characterised how visual adaptation influences the perception of causality by determining the spatial specificity of adaptation and timecourse of recovery from adaptation. In Experiment 1, observers saw ambiguous test events (i.e., the overlap between the two objects varied over trials) at three different horizontal eccentricities. Adaptation was strongest when adaptor and test event were presented at the same eccentricity, and absent when the two were separated by just three degrees of visual angle. Moreover, the perception of causality gradually recovered from adaptation, but remained incomplete. In Experiment 2, both long and short adaptation sequences were highly effective in driving adaptation, and showed no difference in the recovery timecourse, which was complete following more experimental blocks. In Experiment 3, a break without any task-relevant visual input also led to a recovery over the same timespan, but this time, the recovery was instantaneous and incomplete. Altogether, our results provide evidence for highly spatially specific computations, instananeously responding to the onset of adaptation and then gradually recovering from the adaptation over a short time window.

## Introduction

Perception adapts quickly to continuous features in the visual environment—a process known as visual adaptation, in which prolonged exposure to a stimulus temporarily alters subsequent perception. If we stare at the downward motion of a waterfall, and then look at a stationary surface, it appears to move upwards (hence, the waterfall illusion; Addams, 1843). And when we put on sunglasses, the world initially appears tinted by the lenses’ color, but this tint gradually fades over time (de la Hire, 1694; Tregillus & Engel, 2019). These effects typically do not last long, as our system is continuously re-calibrating to the current state of the world. There are, of course, rare exceptions to this general pattern, most famoulsy the McCollough effect, which can persist for days or even months (Jones and Holding, 1975).

Strikingly, visual adaptation extends even to seemingly more high-level visual features, including causal interactions (**Figure 1a**): repeatedly looking at one disc approaching a second disc, setting the second disc into motion, renders later amibiguous test events perceptually less causal (Kominsky & Scholl, 2020; Kominsky & Wenig, 2025; Ohl & Rolfs, 2025; Rolfs et al., 2013). Repeated exposure to one object triggering another’s—where the second motion is considerably faster than the first—results in similar negative aftereffects (Kominsky et al., 2017). These results have supported the view that we understand causal interactions automatically through perceptual processes (Michotte, 1963; Scholl and Tremoulet, 2000) rather than by cognitive reasoning (Rips, 2011; Weir, 1978; White, 2006). This poses questions about the specificity of such adaptation effects in space and time. How does our visual system recover from being adapted to causal interactions, and how spatially precise are the effects of visual adaptation to causal interactions to begin with?

**Figure 1.**
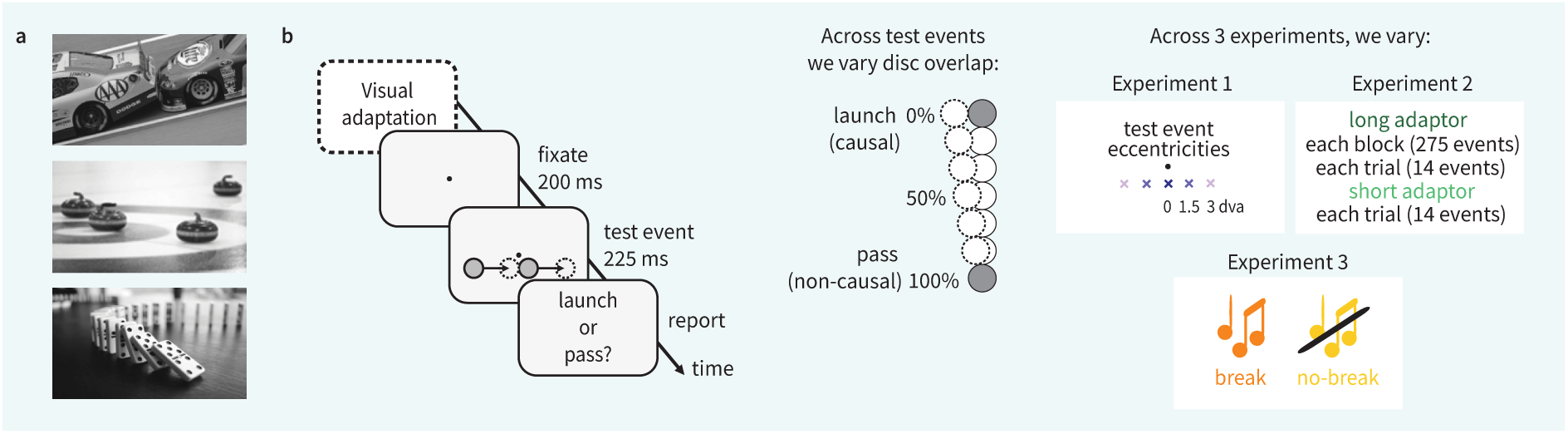
Theoretical background and experimental protocol. **(a)** Examples of causal interactions. **(b)** Basic experimental paradigm, varied across three experiments. In all three experiments, we varied the disc overlap in test events from launch (causal, 0% overlap) to pass (non-causal, 100% overlap) stimuli in 7 equidistant steps. In Experiment 1, we varied the eccentricity of the test event relative to the adaptor. In Experiment 2, we varied the adaptor type (long, short, or non-causal control). In Experiment 3, we introduced a break between adaptation and post-adaptation blocks.

In the present study, we build on the notion that perceiving causal interactions is a perceptual, automatic process that originates in the visual system (Scholl & Tremoulet, 2000). Visual adaptation is a powerful behavioral tool for probing the mechanisms of perception (Kohn, 2007; Webster, 2015), which provides a testing ground for the processes underlying causal perception. More specifically, using visual adaptation we can present a specific adaptor to observers and test whether adaptation transfers to a test event that differs in some aspect from the adaptor. Depending on whether an adaptation transfer is observed, we can then conclude if this particular aspect of the adaptor is critical for the perceptual mechanism of interest or not. Using this approach, visual adaptation to causal events has been shown to operate in a retinotopic reference frame, that is, it is tied to specific locations relative to the fovea (Kominsky & Scholl, 2020; Kominsky & Wenig, 2025; Rolfs et al., 2013). This spatial specificity demonstrates that early, retinotopically organized visual areas are involved in perceiving causal relations. Further evidence for an automatic process of perceiving causal interactions comes from the lack of transfer of visual adaptation across motion directions (Ohl & Rolfs, 2025), suggesting that causality perception relies on direction-selective channels in the visual system. Taken together, such findings support the notion that causality is not inferred solely through higher-level reasoning, but is already decoded at early stages of visual processing.

The suggestion that the perception of causality is the result of low-level visual processes raises the possibility that visual routines underlying the perception of causality are highly spatially specific to a small region of the visual field. The size of receptive fields increases for later processing stages in higher order visual areas (Wandell et al., 2007; Freeman & Simoncelli, 2011). Consequently, a rather broad spatial tuning could falsify the notion that causality perception is an early visual process. One way to address this question is by examining the size of the adapted field. If neurons involved in detecting causal interactions have large receptive fields, adaptation should generalize across a wider area of the visual field. Alternatively, if the receptive fields of these neurons are small, adaptation should be tightly confined to the spatial location of the adapting stimulus. Examining the size of the adapted field thus allows us to better understand where within the visual hierarchy causal perception manifests.

Another aspect of causal perception that remains poorly understood is the temporal dynamics of its underlying visual mechanisms. In other words, after adapting to causal interactions, does perception completely return to baseline, or do residual effects persist? Answering these questions would reveal what kind of information the visual system tracks and the timescales over which it calibrates (Webster, 2015). Given that both the environment and the internal state of the observer vary across multiple timescales, it is reasonable that adaptation mechanisms should also operate with different temporal profiles (Kording et al., 2007). One possibility is that the visual system responds only briefly to causal interactions, producing rapid effects that quickly decay. Such short-lived adaptation would suggest a flexible and responsive mechanism, capable of rapidly adjusting to moment-to-moment changes. Alternatively, adaptation might persist over longer durations, reflecting a more stable recalibration of causal perception.

An additional open question concerns the timecourse of the recovery from adaptation: does the visual system return to baseline immediately, in a step-like fashion, or does it recover slowly over time? While there are many findings on the onset of various visual adaptation effects, there is far less research on how the system recovers from them. One exception is the phenomenon of color afterimages. After fixating on a colored patterns, observers initually perceive a complementary-colored afterimage when the stimulus is removed, but this percept gradually fades and the visual system returns to its baseline state (Zaidi et al., 2012). In motor learning, recovery dynamics have been instrumental for modeling adaptive processes, prompting the development of targeted protocols within the oculomotor domain (Cassanello et al., 2016). Despite this, the timecourse of the recovery from visual adaptation remains largely understudied. Understanding the recovery profile can give insights into how enduring the visual aftereffects of causal adaptation are, and whether they reflect short-term recalibrations or longer-lasting perceptual shifts.

The current experiments had three objectives. First, we assessed how far away from the spatial location of visual adaptation the perception of causal interactions is still affected. Second, we asked how strong of an adaptor we need to observe a shift in the perception of causality, and how quickly this effect might recover back to the starting point. Finally, we asked whether recovery from visual adaptation occurs in a task-relevant manner, through active recalibration of the visual system to the statistical regularities of the environment, or in a time-based manner, resembling a more passive form of recalibration.

To realise this, we made use of the phenomenon that observers readily perceive causality in simple kinematic displays; a moving disc that stops next to another disc appears to launch the second disc into motion (the *launching effect*, **Figure 1b**; Michotte, 1963; see Rips, 2011; Scholl and Tremoulet, 2000; White, 2006, White, 2017 for reviews). To test for spatial specificity of the perception of causal launches, we varied the center point of test events—while keeping all other characteristics constant—and determined whether visual adaptation to launches transfers to test events with a different center eccentricity. If the strength of the adaptation’s perceptual consequences depends on the spatial location of the adaptor and the test event, it would reveal that the mechanisms for the detection of causal interactions are tightly tuned to specific spatial locations. Alternatively, if we observe adaptation transfer from one eccentricity to another, this will support the notion of a broadly tuned mechanism.

To address how the perception of causality recovers from visual adaptation, we presented test events not only before and during, but also after adaptation, once we had removed the adapting stimulus. This design allowed us to assess whether the perception of causality quickly returned to baseline, which would suggest a highly flexible and rapid visual mechanism for the detection of causal interactions. In three experiments, we assessed how spatially specific visual adaptation is, whether the perception of causality completely recovers from adaptation, and whether recovery is task-based or time-based. To preview our results: we provide compelling evidence for highly spatially specific computations underlying the perception of causality, instantaneously responding to the onset of adaptation and then also quickly recovering from visual adaptation in a time-based manner.

## Methods

### Observers

We tested 12 human observers in **Experiment 1** (ages 22–34 y; six female; six male; twelve right-handed; six right-eye dominant), 14 in **Experiment 2** (ages 18–34 y; 7 female; 7 male; 10 right-handed; 10 right-eye dominant), and 12 in **Experiment 3** (ages 18-26 y; 9 female, 3 male; 10 right-handed; 5 right-eye dominant). There were four sessions in **Experiment 1** and **2** (one training session without adaptation, three test sessions with adaptation), and three in **Experiment 3** (one training session, two test sessions with adaptation). The current sample size was comparable with previous work investigating the perception of causality (e.g., Ohl & Rolfs, 2025; Sommer et al., 2025). Based on the data obtained in the session without adaptation (i.e., the first session), we assessed whether observers can distinguish between passes and launches in the basic experiment by identifying whether the proportion of reported passes increases with increasing disc overlap. From **Experiment 2**, two observers were excluded from the final analyses based on this criterion. Data from the first session (that is, without adaptation) did not enter the final analyses. We paid observers €10 per session as compensation for participation. Students enrolled in a Psychology bachelor’s programme could opt for course credits or a combination of money and credits as compensation. We obtained observers’ written informed consent before the first session. All observers had normal or corrected-to-normal vision. The study was approved by the ethics committee of the Psychology Department of the Humboldt-Universität zu Berlin and followed the guidelines of the Declaration of Helsinki (‘World Medical Association Declaration of Helsinki’, 2013).

### Material

Observers sat in a sound- and brightness-attenuated room putting their head on a chinrest. We controlled for observers’ eye position by tracking their dominant eye using an Eyelink 1000+ Desktop Mount eye tracker (SR Research, Ottawa, ON, Canada) with a sampling rate of 1000 Hz. The eye tracker was calibrated (9 points) at the beginning of a session, after breaks, and whenever necessary throughout the experiment. We displayed visual stimuli on a video-projection screen (Celexon HomeCinema, Tharston, Norwich, UK) using a PROPixx DLP projector (VPixx Technologies Inc, Saint Bruno, QC, Canada) at a spatial resolution of 1920×1080 pixels and a refresh rate of 120 Hz. The screen was mounted on a wall at 180 cm away from the observer. The experiment was run on a DELL Precision T7810 (Debian GNU Linux 8) and implemented in Matlab 2023b (Mathworks, Natick, MA, USA) using the Psychophysics toolbox 3 (Brainard, 1997; Kleiner et al., 2007; Pelli, 1997) for stimulus presentation and the Eyelink Toolbox (Cornelissen et al., 2002) for control of the eye tracker. Behavioral responses were collected by registering one of the two possible keypresses on a standard keyboard. In **Experiment 3**, an Apple iPod Nano (5th Generation; Apple Inc., released 2008) with standard wired in-ear headphones was used to present audio.

### General procedure

In all experiments, we asked observers to report whether they perceived a launch or a pass in test events composed of two discs. In the test events, a peripheral disc approached a stationary second disc and stopped at varying disc overlaps across trials (**Figure 1b**). Immediately after that, the second disc started moving in the same direction and with the same speed as the first disc. Before the first trial, observers read the instructions that were displayed on the screen. As part of the instructions, we presented demo trials of test events with 0% overlap that are typically perceived as launches and test events with 100% disc overlap that are typically perceived as passes. Observers had the opportunity to inspect these two events as often as they wanted. Following the instructions, we ran a short training block of 35 (**Experiment 1**) or 28 trials (**Experiment 2 and 3**) with 7 different disc overlaps ranging from 0 to 100% overlap in equidistant steps.

At the beginning of a trial, we asked observers to fixate a gray fixation point with a diameter of 0.2 degrees of visual angle (dva) in the center of the screen on a black background. Trials started once observers successfully fixated for at least 200 ms. We presented the test events at 3 dva below the fixation point. In **Experiment 1**, the center of the test event varied between 0, ±1.5, and ±3 dva along the horizontal axis (relative to the fixation point. In test events, the first moving disc (gray; diameter of 1.5 dva) located either left or right from the vertical meridian started moving towards a stationary disc (gray; diameter of 1.5 dva). The first disc stopped moving at one of seven possible distances away from the stationary disc, resulting in 7 different disc overlaps ranging from 0 to 100% in equidistant steps. The entire duration of the test event was 225 ms. At the end of the trial, observers reported whether they perceived a launch or a pass in the test event by pressing either the arrow up key or the arrow down key.

In addition, the first session served to determine whether observers perceived launches and passes as a function of disc overlap. To this end, in **Experiment 1**, observers completed a training session with 10 blocks in which we varied the horizontal center point of the test events (within-subjects: 5 horizontal center eccentricities at 0, ±1.5, and ±3 dva from the fixation point) and the amount of disc overlap (within-subjects: 7 disc overlaps ranging from 0 to 100% overlap, in equidistant steps). We presented these conditions in a randomized order within a block. Each combination of these manipulations was presented once in a block, resulting in 35 trials in each block and a total of 350 trials in the first session. In **Experiments 2** and **3**, observers also completed 10 blocks in the first session, but the horizontal center point of the test events was always at 0 dva from the fixation point. Instead, every combination of the disc overlap manipulation was repeated four times in each block, resulting in 28 trials in a block and a total of 280 trials in the first session. In all experiments, we counterbalanced the direction of the adaptation and test event (left-to-right vs. right-to-left) and the mapping of the response keys to reflect ‘launch’ or ‘pass’ (arrow up vs. arrow down key) between observers. In sessions 2–4 of **Experiment 2**, we presented three different adaptor types that varied between the experiments. In sessions 2 and 3 of **Experiment 3**, we introduced a break between adaptation and post-adaptation blocks in either of the two sessions. See details of the procedures for the specific experiments below.

In all experiments, we tracked observers’ dominant eye to ensure proper fixation behavior during presentation of the test events and presentation of the adaptors. More specifically, we tracked the dominant eye’s current position at a sampling rate of 1000 Hz and determined the eyes’ deviation from the fixation point online. We aborted a trial whenever this distance exceeded 2 dva. Observers repeated these trials at the end of a block in randomized order. During presentation of the adaptors, we presented a short message (at the fixation point) saying “please fixate” if their gaze exceeded 2 dva away from the fixation point.

### Adaptation in Experiment 1

In **Experiment 1**, each observer completed 18 blocks in each of sessions 2–4. The first six blocks in a session were without adaptation to measure an observer’s perception of causality before adaptation. As in the first session, we presented test events at five possible horizontal center eccentricities and varied the disc overlap, resulting in 35 trials in each block. In blocks 6–12, we presented an adaptor before the first trial of a block. The adaptor was a stream of launching events in the same direction as the test event. To this end, we presented 275 launching events before the first trial of a block, each located at the same horizontal offset as the fixation point (0 dva) and vertically at 3 dva below the fixation point. The exact direction of these launching events were randomly chosen from a narrow uniform distribution around the direction on the horizontal meridian (±30 degrees) to avoid adaptation to low-level features (such as contrast and luminance) rather than specifically to the causal percept (Ohl & Rolfs, 2025). The long adaptation phase was complemented by a top-up adaptation of 14 launching events in the same direction as the adaptor (drawn from the same narrow uniform distribution around the main direction) before each trial to keep adaptation effective throughout the entire block. Blocks 13–18 were without adaptation, so that we could address the recovery from visual adaptation. Each observer completed a total of 1,890 trials in **Experiment 1**.

### Variations in Experiment 2

In **Experiment 2**, we presented three different adaptor types to address the strength of the recovery of visual adaptation. Each of sessions 2–4 consisted of a different type of adaptor: one which was exactly as in **Experiment 1,** which combined the long adaptation sequence of 275 launching events at the start of each adaptation block with 14 top-up events in each adaptation trial (‘long’ adaptation), one which only had the 14 top-up events (‘short’ adaptation) and a non-causal control. The purpose of the short adaptor was to see whether it would elicit the same adaptation response as a longer adaptor. For the control condition, we used non-causal ‘slip’ events (Rolfs et al., 2013) as adaptors to control that the adaptation was specific for the impression of causality in the launching events. These control events matched launching events in as many physical properties as possible while producing a very different, non-causal phenomenology. In non-causal control events, the first disc also moved towards a stationary second disc. In contrast to launching events, however, the first disc passed the stationary disc and stopped only when it was adjacent to the opposite edge of the stationary disc. While slip events do not elicit a causal impression, they have the same number of objects and motion onsets, the same motion direction and speed, as well as the same spatial area of the event as launches. As for the launch adaptors, we displayed control adaptors in a narrow uniform range of directions around one of the two possible horizontal directions (from left-to-right or from right-to-left being fixed per observer). This experimental design resulted in three test sessions of which the order was randomly determined for each observer. This time, blocks 13–22 were without adaptation, so that we could address the recovery from visual adaptation over a longer time interval as compared to **Experiment 1** (6 blocks of post-adaptation in **Experiment 1** vs. 10 post-adaptation blocks in **Experiment 2**). Each observer completed a total of 1,848 trials in **Experiment 2**.

### Variations in Experiment 3

In **Experiment 3**, we presented only top-up adaptors, as our results from **Experiment 2** suggested that short initial adaptation events sufficed to elicit the same magnitude of adaptation as a long initial adaptation event. In **Experiment 3**, we aimed to address whether we recover from visual adaptation in a task-relevant manner by actively recalibrating the visual system to the statistical regularities in the environment, or whether this is a more passive, time-based process. That is, do we recover from adaptation by seeing numerous non-causal events in the environment? To test this, we introduced a break without task-relevant visual input after the adaptation blocks (after block 12). During this break, observers were required to stay seated and listen to one of three podcast options: “The Science of Birth Without Pregnancy” (O’Neill, 2021), “The Science of Designer Babies” (O’Neill, 2024a), or “The Science of Heart Attacks” (O’Neill, 2024b). The purpose of the podcast was to present entertainment whilst simultaneously not presenting any visual causal interactions of any kind. This allowed us to address whether the perception of causality also recovered when there was no task-relevant input that would suggest different statistical regularities of the environment, while time did pass. The podcast was exactly 10 minutes long, matching the average time the post-adaptation blocks took in **Experiment 2**. The observer’s eyes were not tracked during the podcast break. This experimental design resulted in two test sessions of which we counterbalanced the order for each observer. Like in **Experiment 2**, blocks 13–22 were without adaptation, so that we could address the recovery from visual adaptation over a longer time interval compared to **Experiment 1** (i.e., 10 rather than 6 post-adaptation blocks). Each observer completed a total of 1,848 trials in **Experiment 3**.

### Data analysis

For the statistical analyses and estimation of the psychometric functions, we used the *quickpsy* package (Linares, 2016) and the RStudio environment (Posit team, 2025). We related disc overlap to the proportion of reported launches using logistic functions with four parameters for the intercept, slope, as well as upper and lower asymptotes. We fitted these functions separately for each observer and condition and obtained the points of subjective equality (PSE; the amount of overlap between the two discs in the test events that result in the same proportion of reported launches and passes). For model fitting, we constrained the range of possible estimates for each parameter of the logistic model. The lower asymptote for the proportion of reported launches was constrained to be in the range of 0–0.75, and the upper asymptote in the range of 0.25–1. The intercept of the logistic model was constrained to be in the range 1–15, and the slope was constrained to be in the range –20 to –1.

To investigate patterns of the perception of causality block-by-block and over the different phases of the experimental sessions (before, during, and after adaptation), we related causal reports over blocks and phases to the three different testing eccentricities (0 dva = center; ±1.5 dva = near; ±3 dva = far) in **Experiment 1**, the three different adaptors (long, short, control) in **Experiment 2**, and the two break conditions (podcast-break or no-break) in **Experiment 3**. Again, we calculated the mean proportions separately for each observer and condition and then calculated group means. To confirm that symmetrical eccentricities could be collapsed in **Experiment 1**, we statistically compared the psychometric thresholds at each eccentricity (−1.5 vs +1.5 dva; −3 vs +3 dva) using paired t-tests. No significant differences were found for either the near or far conditions in any phase (all *p*s ≥ .19), indicating that symmetric positions produced equivalent PSEs. Therefore, we collapsed ±1.5 dva into a single “near” level and ±3 dva into a “far” level.

For inferential statistics, we analyzed PSEs using repeated-measures analyses of variance (rmANOVA). Error bars indicate Cousineau–Morey 95% confidence intervals (Morey, 2008). A significant interaction in the rmANOVA was complemented by running post-hoc paired t-tests.

The model comparisons for non-linear mixed effects models were based on Akaike and Bayesian Information Criteria (AIC, BIC) computed from maximum-likelihood fits.

**Experiments 1** and **3** were preregistered at the Open Science Framework (https://osf.io/e4yjn/).

## Results

In three experiments, we tested how the perception of causality recovers from visual adaptation. Previous adaptation studies on causality perception have typically compared the proportion of launches (or points of subjective equality, PSEs) before adaptation to those measured during adaptation. To our knowledge, no study has additionally compared the proportion of launches after adaptation to the pre-adaptation and adaptation phases. Here, we evaluated recovery by directly contrasting the PSEs after adaptation to those measured before and during adaptation.

### Adaptation to causal events is spatially specific

In **Experiment 1**, we examined the points of subjective equality (PSEs) across the three phases (pre, adaptation, post) as a function of test event eccentricity (center, near, far) using a two-way repeated-measures ANOVA, followed by pairwise contrasts. Moreover, we fitted non-linear mixed effects models designed to answer whether there was an adaptation effect (pre-adaptation vs. adaptation), if so, whether recovery occurred, (post-adaptation vs. adaptation), and if so, whether the recovery was complete (post-adaptation vs. pre-adaptation). We additionally tested whether adaptation to launch events is spatially specific to the center of the adapting stimulus, or whether it spreads to nearby test event eccentricities. To this end, we presented test events at one of three possible horizontal eccentricities from the fixation point (0, ±1.5, and ±3 dva; **Figure 2a**). The first disc of the test event stopped as soon as the two partially overlapped (manipulated in 7 equidistant steps from no to full overlap). The second disc then started to move in the same direction as the first disc, and observers reported whether they perceived that the first disc launched the second (common at no overlap) or passed it (common at full overlap). We quantified the PSE between perceived launches and passes by modeling psychometric functions that related disc overlap to the proportion of launch reports before, during, and after adaptation.

**Figure 2.**
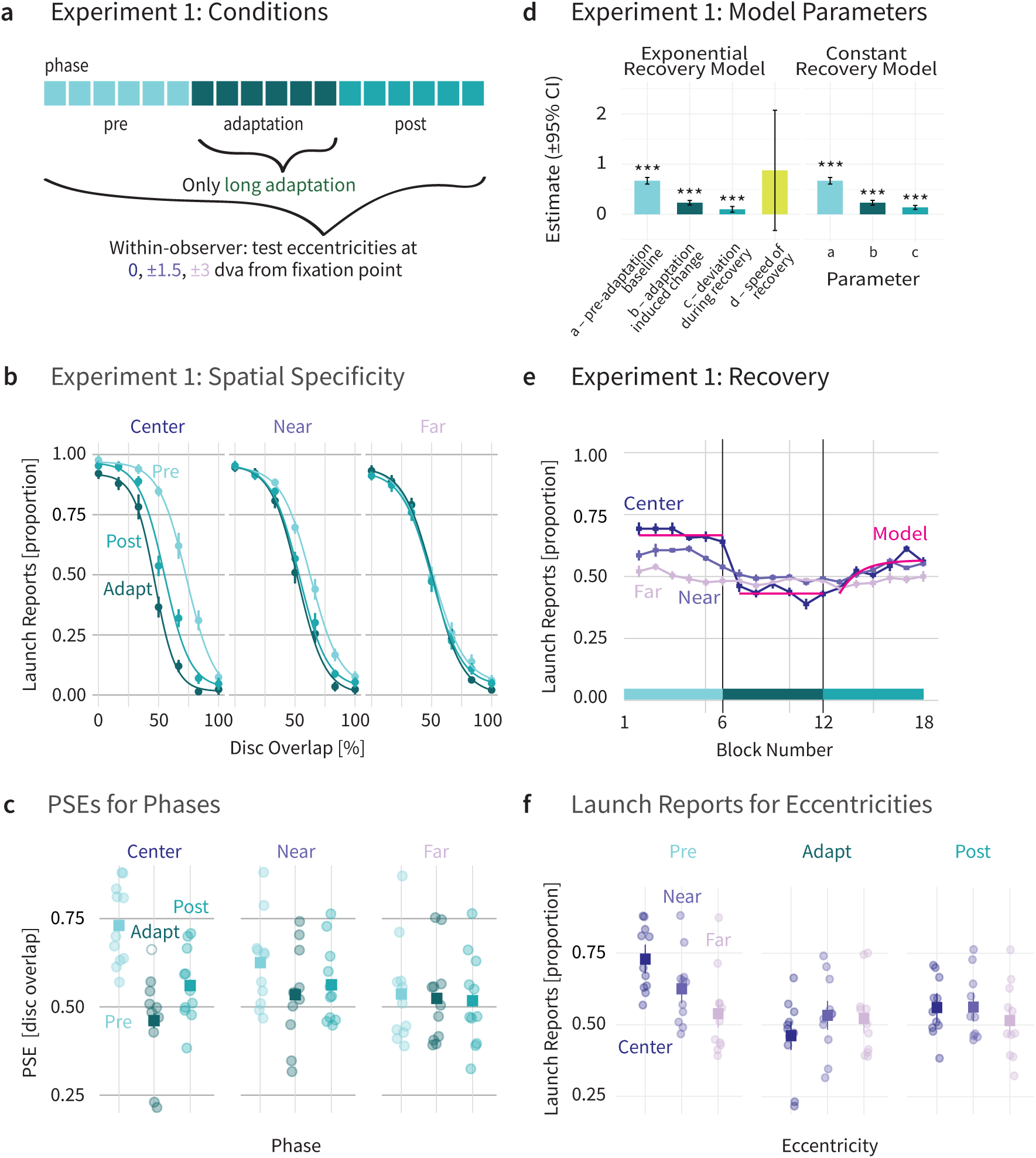
Results of Experiment 1. **(a)** Experiment 1 consisted of 18 blocks, divided into pre-adaptation, adaptation, and post-adaptation phases. The adaptor consisted of 275 events at the start of each block, and 14 events at the start of each trial. Test events were presented at horizontal eccentricities at 0, ±1.5, and ±3 dva from the fixation point. **(b)** Mean proportion of launch reports as a function of disc overlap. Visualization of psychometric curves is based on the mean parameters averaged across observers before adaptation (light blue), during adaptation (dark blue), and after adaptation (mid blue) for the different test event eccentricities: center (dark purple), near (mid purple) and far (light purple). **(c)** PSEs for individual observers (circles, n = 12) and the mean across observers (square) for the different test event eccentricities. **(d)** Estimated parameters for competing exponential and constant piecewise non-linear mixed effects models. Model fitting was restricted to trials at the center eccentricity (0 dva), as this was the only eccentricity showing reliable adaptation and recovery effects. Parameter a denotes the pre-adaptation baseline, b denotes the adaptation-induced change relative to the baseline, c denotes the asymptotic deviation from baseline during recovery, and d denotes the speed of the recovery. For each of the parameters, we denote 95% confidence intervals. *p < .05, **p < .01, ***p < .001. **(e)** Blockwise analysis of mean proportion of launch reports. Visualization of blockwise averages is based on the proportion of launch reports per block in the experiment, averaged across observers for each test event eccentricity. Additionally, the winning non-linear mixed effects model fit (exponential model, dark pink) for trials at the center eccentricity (0 dva). **(f)** Launch reports for individual observers (circles) and the mean across observers (square) for the block numbers. Error bars indicate Cousineau–Morey 95% confidence intervals (Morey, 2008).

Visual adaptation reduced the perception of causality at the center and near test event eccentricities, but not for the far eccentricity (**Figure 2b,c**). Observers reported fewer launches during adaptation compared with pre-adaptation, with the strongest effect at the center (PSE_pre_ 0.675±0.041; PSE_adaptation_ = 0.444±0.049; ΔPSE = –0.231±0.049), weaker effects at the near eccentricity (PSE_pre_ = 0.591±0.031; PSE_adaptation_ = 0.500±0.032; ΔPSE = –0.091±0.031), and no adaptation at the far eccentricity (PSE_pre_ = 0.508±0.045; PSE_adaptation_ = 0.486±0.024; ΔPSE = –0.022±0.024). After adaptation, PSEs partially recovered during the post-adaptation phase (**Figure 2e**; center: PSE_post_ = 0.538±0.033; ΔPSE = –0.137±0.033; near: PSE_post_ = 0.532±0.019; ΔPSE = –0.059±0.019; far: PSE_post_ = 0.486±0.046; ΔPSE = –0.022±0.046). A two-way rmANOVA (3 phases: pre-adaptation, adaptation, post-adaptation; 3 eccentricities: center, near, far) on the PSEs revealed significant main effects of phase (F(2, 22) = 16.23, p < .001) and eccentricity (F(2, 22) = 5.997, p = .008), as well as a significant interaction between phase and eccentricity (F(4, 44) = 32.80, p < .001).

A one-way rmANOVA on pre-adaptation PSEs revealed a significant effect of eccentricity (F(2, 22) = 20.20, p < .01) already in the pre-adaptation phase. Pairwise comparisons showed that PSEs were higher at the far eccentricity compared with both near (Δ = 0.083, p < .001) and center (Δ = 0.167, p < .001) positions, and higher at near compared with center (Δ = 0.084, p < .001), indicating that baseline causal perception varies systematically with test event eccentricity.

Estimated marginal mean contrasts confirmed the spatial specificity of the adaptation effect. First, pre vs. adaptation (testing the adaptation effect) showed a large reduction at the center (estimate = 0.208±0.039, p < .001), smaller but reliable effects at the near eccentricity (estimate = 0.068±0.037, p = < .001), and no effect at the far eccentricity (estimate = – 0.002±0.037, p = 1.000). Pairwise contrasts further showed that the adaptation effect at the center was larger than at near and far, and near was larger than far (p < .05). Second, post vs. adaptation (testing recovery) revealed significant recovery at the center (estimate = – 0.084±0.039, p < .001), but not at the near (estimate = –0.022±0.037, p = .247) or far eccentricity (estimate = 0.011±0.037, p = .571).

### Recovery from adaptation is rapid but gradual

To characterize the temporal dynamics of the recovery from adaptation in more detail, we modeled responses across blocks using a piecewise non-linear mixed-effects model. This model consisted of three pieces: a constant pre-adaptation baseline, a constant adaptation level, and an exponential recovery function during the post-adaptation phase. Responses 𝑦(𝑥) as a function of block number 𝑥 were modeled as

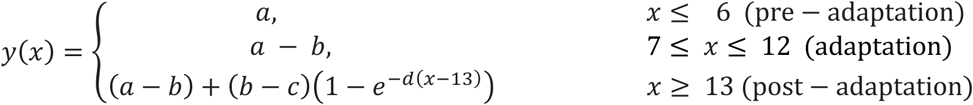

where 𝑎 denotes the pre-adaptation baseline, 𝑏 denotes the adaptation-induced change relative to the baseline, 𝑐 denotes the asymptotic deviation from baseline during recovery, and 𝑑 denotes the speed of the recovery. This model, assuming a gradual recovery via the exponential function, we then compared to a reduced model assuming an instantaneous, step-like recovery. This reduced model omitted the exponential term and assumed a constant post-adaptation level 𝑎 − 𝑐. Random effects were included on parameters 𝑎, 𝑏, and 𝑐 to account for between-observer variability. Both models were restricted to trials at the center eccentricity, as this was the only eccentricity showing reliable adaptation and recovery effects; near and far eccentricities were therefore excluded from this analysis.

Model comparison favored the exponential model, which showed lower information criteria (AIC = 5234.26, BIC = 5302.97) than the reduced constant recovery model (AIC = 5241.44, BIC = 5303.90). While BIC showed only a marginal preference, AIC provided support for gradual recovery dynamics. This indicates that recovery unfolded progressively over multiple post-adaptation blocks rather than occurring abruptly at the transition from adaptation to post-adaptation (**Figure 2d–f**). Together, these results demonstrate that recovery from adaptation was gradual in **Experiment 1**. Importantly, despite this gradual recovery, post-adaptation did not fully return to pre-adaptation levels. Parameter 𝑐, the asymptotic deviation from baseline during recovery, was significantly greater than zero (𝑐 = 0.099, SE = 0.029, p < .001). Thus, the observed recovery from adaptation was incomplete in this experiment.

### Recovery from adaptation is complete

In **Experiment 2,** we extended the post-adaptation phase compared to **Experiment 1** to assess whether the perception of causality fully recovers with more time. Observers experienced three adaptor types: long, short, and a non-causal control (**Figure 3a**). Points of subjective equality (PSEs) were quantified before, during, and after adaptation for the three adaptor types. As before, we tested whether adaptation occurred (pre vs. adaptation), whether recovery occurred (post vs. adaptation), and whether recovery was complete (post vs. pre).

**Figure 3.**
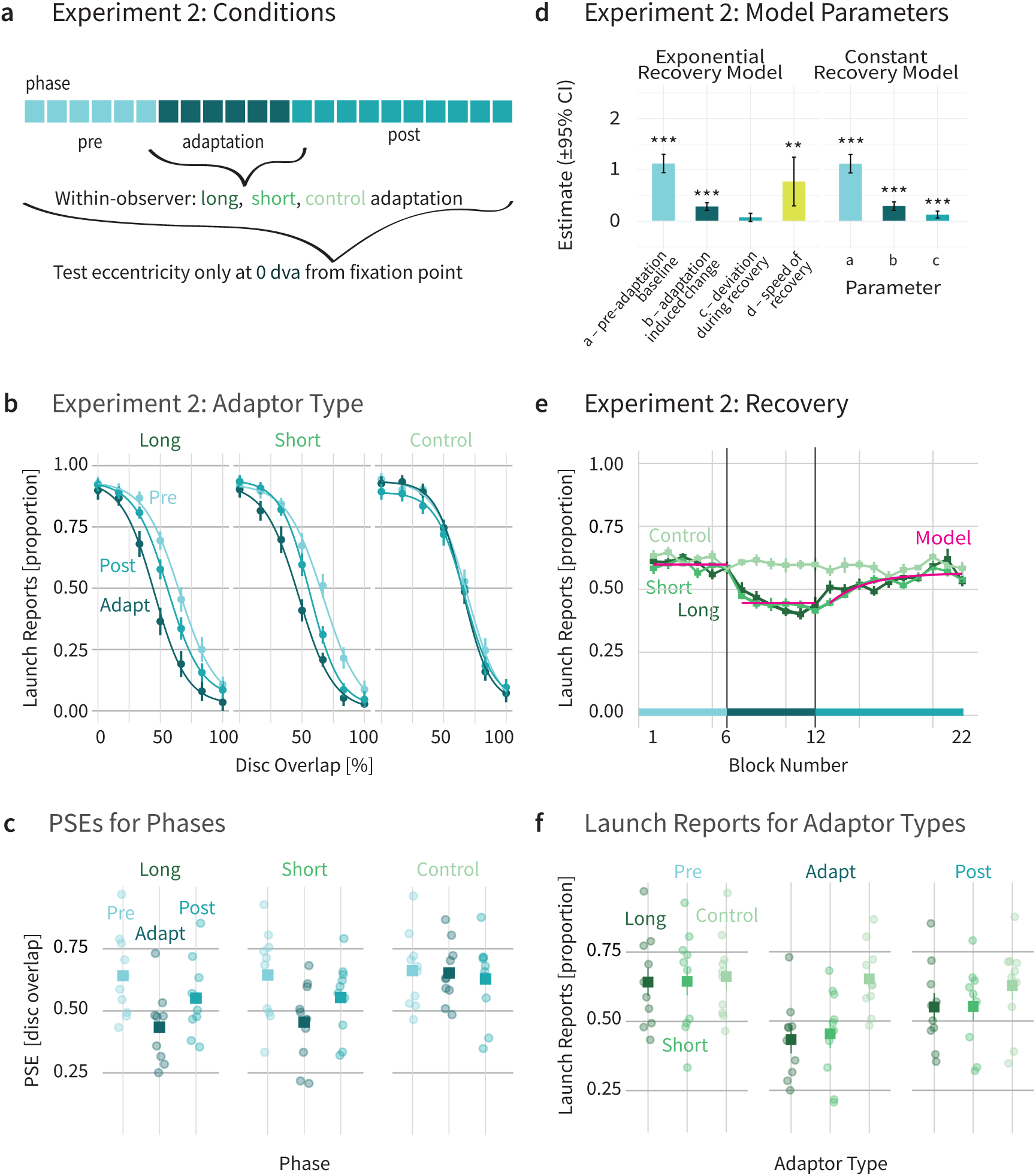
Results of experiment 2. **(a)** Experiment 2 consisted of 22 blocks, divided into pre-adaptation (6), adaptation (6), and post-adaptation (10) phases. The long as well as the control adaptor consisted of 275 events at the start of each block, and 14 events at the start of each trial. The short adaptor consisted only of 14 events at the start of each trial. **(b)** Mean proportion of launch reports as a function of disc overlap. Visualization of psychometric curves is based on the mean parameters averaged across observers before adaptation (light blue), during adaptation (dark blue), and after adaptation (mid blue) for the different adaptor types: long (dark green), short (mid green), and the non-causal control (light green). **(c)** PSEs for individual observers (circles, n = 12) and the mean across observers (square) for the different adaptor types (same order as in the above panel). **(d)** Estimated parameters for competing exponential and constant piecewise non-linear mixed effects models. For model fitting, trials from long and short adaptors were collapsed, since we observed no difference between the two adaptors. Parameter a denotes the pre-adaptation baseline, b denotes the adaptation-induced change relative to the baseline, c denotes the asymptotic deviation from baseline during recovery, and d denotes the speed of the recovery. For each of the parameters, we denote 95% confidence intervals. *p < .05, **p < .01, ***p < .001. **(e)** Blockwise analysis of mean proportion of launch reports. Visualization of blockwise averages is based on the mean launch reports per block in the experiment, averaged across observers for the long adaptor (dark green), short adaptor (mid green) and non-causal control adaptor (light green). Additionally, the winning non-linear mixed effects model fit (exponential model, dark pink) for trials from long and short adaptors. **(f)** Launch reports for individual observers (circles, n = 12) and the mean across observers (square) for the different phases. In all subfigures, error bars indicate Cousineau–Morey 95% confidence intervals (Morey, 2008).

During the adaptation phase, long and short adaptors produced clear decreases in causal perception (**Figure 3b,c**; long: ΔPSE = –0.154±0.124; short: ΔPSE = –0.147±0.109), whereas the non-causal control adaptor produced no meaningful change (ΔPSE = – 0.015±0.105). In the post-adaptation phase, causal perception partially recovered for long and short adaptors (**Figure 3e**; long ΔPSE = –0.061±0.104; short ΔPSE = –0.063±0.088), while the control remained near baseline (ΔPSE –0.031±0.118).

A two-way repeated-measures ANOVA with factors phase (pre, adaptation, post) and adaptor type (long, short, control) revealed significant main effects of phase (F(1.71, 18.86) = 13.84, p < .001, ges = .065) and adaptor type (F(1.60, 17.55) = 9.58, p = .003, ges = 0.049), as well as a significant interaction (F(2.63, 28.98) = 8.89, p < .001, ges = 0.027), indicating that the magnitude of adaptation depended on the adaptor type.

Planned contrasts testing the adaptation effect (pre vs. adaptation) confirmed significant reductions in PSEs for both the long (estimate = 0.154±0.033, p < .001) and short (estimate = 0.147±0.017, p < .001) adaptors relative to baseline, whereas no adaptation effect was observed for the control adaptor (estimate = –0.084±0.027, p = .649). Direct comparisons showed that adaptation effects were significantly larger for the long and short adaptors than for the control (long vs. control: t(11) = –3.83, p < .001; short vs. control: t(11) = –5.56, p < .001), while the long and short adaptors did not differ from each other (t(11) = 0.02, p = 1.000). To provide additional evidence that the long and short adaptors were indeed not different, we computed a Bayesian paired t-test. This provided moderate evidence for the absence of a difference (BF_01_ = 3.48).

We next assessed the dynamics of the recovery from adaptation by fitting the same two competing piecewise nonlinear mixed-effects models as in **Experiment 1**: a model with an exponential recovery component and a reduced model assuming instantaneous (constant) recovery. Both models were fit after excluding the slip adaptor condition; only the long and short adaptor conditions were included in the modeling. Model comparison revealed a modest advantage for the exponential model, which provided a lower AIC than the constant model (19914.7 vs. 19917.2) (**Figure 3d–f**). However, BIC values were very similar between the two models (19998.4 vs. 19993.3). Therefore, we caution for an overinterpretation of the evidence for the exponential model, noting that while a gradual recovery is plausible, the data do not rule out a simpler constant-recovery account. Finally, to assess whether recovery was complete, we compared post-adaptation reports to pre-adaptation baseline by investigating parameter 𝑐, the asymptotic deviation from baseline during recovery. The parameter was not sigificantly different from zero (𝑐 = 0.033, SE = 0.022, p = .135), suggesting that post-adaptation reports returned to pre-adaptation baseline. Recovery after adaptation was therefore complete in **Experiment 2**.

Together, the results of **Experiment 2** show that adaptation to causal launch events robustly reduces perceived causality, that this reduction weakens once the adaptor is removed, and that extending the post-adaptation phase leads to gradual and complete recovery. Importantly, recovery dynamics did not differ between long and short adaptors, and no adaptation or recovery was observed for a non-causal control adaptor, mirroring the pattern of specificity observed in **Experiment 1**.

### Recovery from adaptation is time-based

In **Experiment 3**, we asked whether the perception of causality recovers in the absence of experimentally controlled task-relevant visual input, that is, purely as a function of time. To this end, observers either took a 10-minute “podcast-break” between the adaptation and post-adaptation phase (viewing a neutral display), or had no break between the two phases (**Figure 4a**). We again quantified PSEs before, during, and after adaptation to assess if there was an adaptation effect (pre vs. adaptation), and if so, whether recovery occurred (post vs. adaptation).

**Figure 4.**
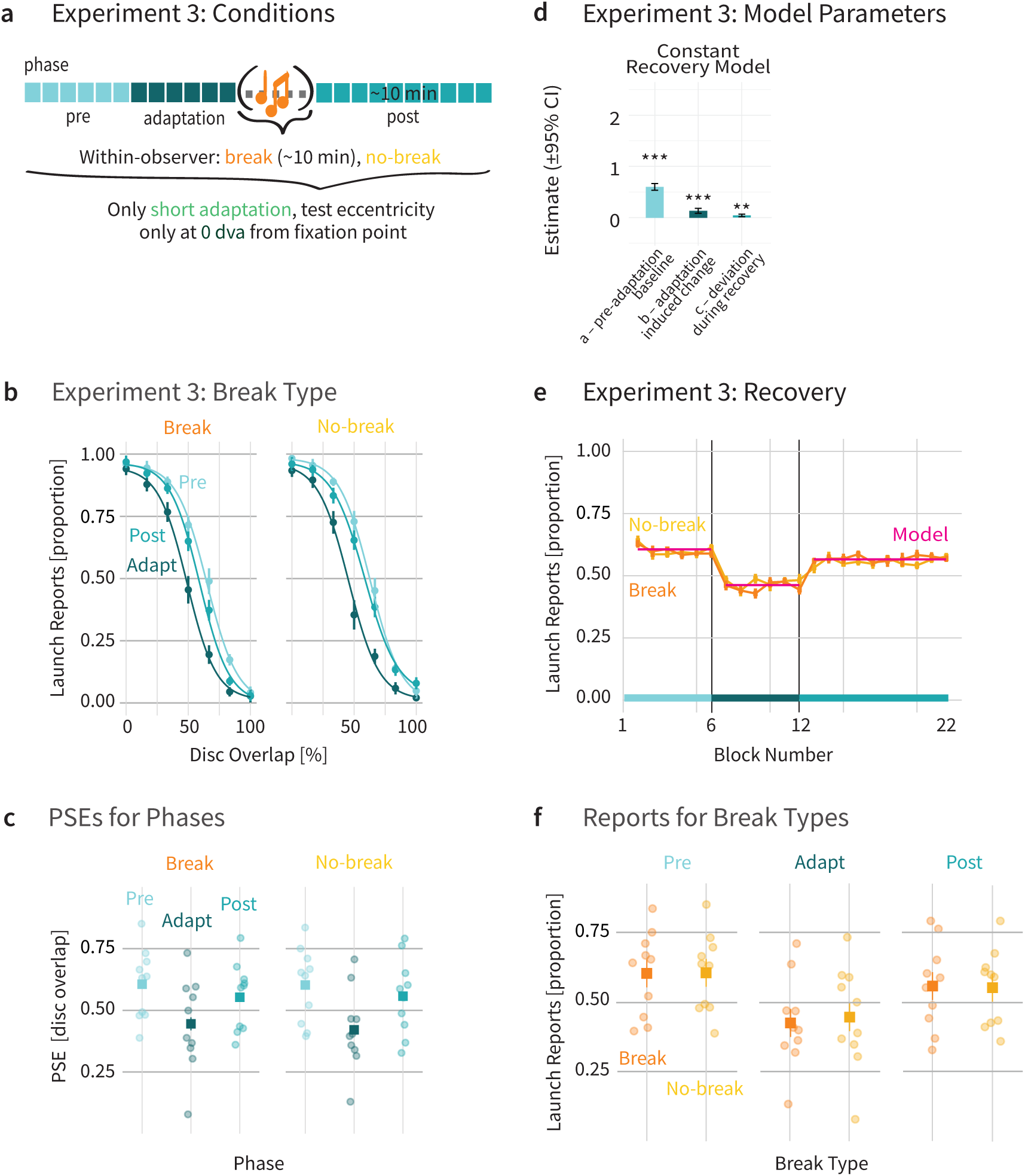
Results of Experiment 3**. (a)** Experiment 3 consisted of 18 blocks, divided into pre-adaptation, adaptation, and post-adaptation phases. The adaptor consisted of 14 events at the start of each trial. Observers took part in two testing sessions, one with a podcast break within the adaptation and post-adaptation phases, and one session without a break. **(b)** Mean proportion of launch reports as a function of disc overlap. Visualization of psychometric curves is based on the mean parameters averaged across observers before adaptation (light blue), during adaptation (dark blue), and after adaptation (mid blue) for the different break conditions: break (orange), and no-break (yellow). **(c)** PSEs for individual observers (circles, n = 12) and the mean across observers (square) for the different break conditions (same order as in the above panel). (**d)** Estimated parameters for the constant piecewise non-linear mixed effects models. Parameter a denotes the pre-adaptation baseline, b denotes the adaptation-induced change relative to the baseline and c denotes the asymptotic deviation from baseline during recovery. For each of the parameters, we denote 95% confidence intervals. *p < .05, **p < .01, ***p < .001. **(e)** Blockwise analysis of mean proportion of launch reports. Visualization of blockwise averages is based on the mean launch reports per block in the experiment, averaged across observers for the break (orange) and no-break conditions (yellow). Additionally, the winning non-linear mixed effects model fit (constant model, dark pink). **(f)** Launch reports for individual observers (circles, n = 12) and the mean across observers (square) for the different phases. In all subfigures, error bars indicate Cousineau–Morey 95% confidence intervals (Morey, 2008).

PSEs were nearly identical across the two break conditions in all three phases (**Figure 4b,c**; pre: PSE_break_ = 0.604±0.082; PSE_no-break_ = 0.604±0.082; adaptation: PSE_break_ = 0.475±0.082; PSE_no-break_ = 0.459±0.082; post: PSE_break_ = 0.559±0.081; PSE_no-break_ = 0.574±0.081). A two-way repeated-measures ANOVA (3 phases: pre, adaptation, post, 2 break types: podcast-break, no-break) revealed a significant main effect of phase (F(2, 22) = 16.74, p < .001), indicating that adaptation reliably reduced PSEs compared to the pre-adaptation phase. Neither the main effect of break type (F(1, 11) = 0, p = 1.000) nor the interaction between phase and break type (F(2, 22) = 1.84, p = .182) was significant, indicating that a 10-minute podcast break did not reliably modulate adaptation or recovery.

As in **Experiment 1** and **Experiment 2,** we modeled the temporal profile of the recovery from adaptation by fitting the same two competing piecewise nonlinear mixed-effects models. Since both break types showed similar adaptation and recovery effects, we collapsed the two conditions for model fitting. As before, we compared a model with an exponential recovery component to a reduced model assuming instantaneous (constant) recovery. In contrast to **Experiment 1** and **Experiment 2,** the exponential model failed to converge for **Experiment 3**, despite multiple starting values and reduced random-effects structures. This failure reflects the absence of detectable curvature in the post-adaptation data: recovery occurred instananeously following adaptation offset and showed no evidence of a gradual, time-dependent progression therafter. The constant model converged reliably (AIC = 20366.4, BIC = 20442.4), consistent with an abrupt reduction in the adaptation effect following adaptation offset (**Figure 4d–f**). Thus, recovery in **Experiment 3** was best characterized as instantaneous rather than gradual. Finally, to assess whether recovery was complete, we compared post-adaptation reports to pre-adaptation baseline by investigating parameter 𝑐, the asymptotic deviation from baseline during recovery. The parameter remained significantly above zero (𝑐 = 0.039, SE = 0.013, p < .001), suggesting that post-adaptation reports did not fully return to baseline. Thus, recovery in **Experiment 3** was instantaneous but incomplete, consistent with a rapid offset of the adaptation effect followed by a persistent residual bias.

Together, these results show that in **Expeirment 3,** the negative aftereffect in perceived causality weakens over time, but recovery remains incomplete even after a 10-minute interruption. Importantly, the temporal break did not facilitate recovery relative to continuous viewing, indicating that decay over time alone was insufficient to fully reverse adaptation to causality.

## Discussion

We set out to investigate the spatial specificity and temporal dynamics of visual adaptation to causal events. Across three experiments, we found that the effects of visual adaptation were narrowly tied to the spatial location of the test event. Moreover, the recovery from adaptation unfolded quickly in a gradual fashion in **Experiment 1** and **2**, and instantaneously in a time-based manner in **Experiment 3**. In all three experiments, causality perception returned towards baseline levels within the same testing session. We replicated the general adaptation effect reported in previous studies (e.g., Kominsky & Scholl, 2020; Kominsky & Wenig, 2025; Ohl & Rolfs, 2025; Rolfs et al., 2013), showing that visual adaptation reliably alters the perception of causality. In **Experiment 1**, we observed a negative aftereffect during adaptation that varied with eccentricity: adaptation effects were strongest at the adapted location and diminished with increasing distance. Moreover, the perception of causality gradually recovered once we removed the adaptor, but was incomplete within the tested timeframe. In **Experiment 2**, we extended this finding by showing that the recovery was complete with slightly more time. Additionally, a shorter adaptation duration was equally effective in driving the adaptation effect. In contrast to the gradual recovery observed in **Experiments 1** and **2, Experiment 3** showed instantaneous recovery that progressed in a time-based manner even without task-relevant input.

The spatial specificity of the observed aftereffects suggests that visual adaptation to causal events might happen at an early stage in the visual hierarchy. Since adaptation was tightly confined to the spatial location of the test event, it seems likely that neurons involved in detecting causal interactions have rather small receptive fields. Small receptive fields are a hallmark of early visual areas such as V1 and V2, where neural tuning is tightly linked to precise spatial locations (Wandell et al., 2007). By contrast, later visual areas typically integrate information across larger regions of space, supporting position-invariant representations. The fact that adaptation here was spatially specific therefore points to an early stage in visual processing, before such integration occurs. This is consistent with the idea that even seemingly high-level constructs like causality may be grounded in low-level perceptual mechanisms (Michotte, 1963; Scholl and Tremoulet, 2000), with specialized neurons encoding the spatial and temporal properties that give rise to causal impressions.

Our finding of high spatial specificity has important implications for how visual adaptation to causality should be interpreted. If the effect were driven by cognitive factors such as a shift in a decision criterion, that is, a change in the internal threshold for judging whether an event is a collision, then this criterion should, in principle, apply globally across the visual field. A collision is a collision, regardless of where it occurs. However, the adaptation effect we observed was *highly* spatially specific, which is difficult to reconcile with a purely criterion-based account. Although we cannot fully rule out the possibility of a spatially specific decision criterion, such a mechanism would be a rather unlikely feature of the visual system. Instead, the strong spatial specificity is more consistent with the idea that our experimental manipulation altered a visual, perceptual representation itself rather than a global decision threshold. In this sense, the spatial pattern of our results provides evidence that the adaptation effect reflects changes within visual processing rather than shifts in higher-level judgment.

An important contribution of the present work is that it highlights the value of examining not only the onset of visual adaptation, but also its recovery. This recovery period offers a largely untapped time window in which to learn about the dynamics of the visual system. In our case, we observed recovery on a relatively fast timescale compared to the more commonly studied long-term examples, yet within our study this recovery was still more gradual than one might expect from such a brief adaptation phase. Notably, we found that the same adaptation and recovery patterns emerged for both a short and a longer adaptation duration, suggesting that these dynamics are robust across different exposure durations. This has practical implications for future paradigms investigating adaptation in the perception of causality: using shorter adaptors may suffice, allowing for faster trials without compromising the effects.

Our focus on recovery allows us to situate our findings within the spectrum of timescales of recovery observed in other adaptive processes. At the milisecond timescale, for instance, recovery from saccadic adaptation (e.g., Cassanello et al., 2016, 2019; McLaughlin, 1967; Rolfs et al., 2010) and color adaptation (e.g., Zaidi et al., 2012) reveals aspects of recalibration that remain invisible when focusing solely on the adaptation phase (for two exceptions, see Ethier et al., 2008; Wong & Shelhamer, 2011). In contrast, some adaptive phenomena recover over much longer timescales: the McCollough effect can persist for days or even months (Jones & Holding, 1975), and chromatic shifts after donning tinted lenses fade gradually, documented as early as de la Hire (1694). Positioned in the middle of this spectrum, our findings highlight how recovery trajectories differ across domains and show how directly studying recovery can improve our understanding of visual processing more broadly.

To our surprise, recovery profiles were inconsistent between the three experiments, varying both in their onset and whether they completely returned to baseline. In **Experiment 1**, recovery unfolded gradually but remained incomplete. In **Experiment 2**, recovery was likewise gradual, yet reached baseline levels, indicating complete recovery. In contrast, **Experiment 3** showed an instananeous onset of recovery following adaptation offset, but this recovery again remained incomplete. One possible explanation for this variability is learning or memory consolidation across the experimental session: repeated exposure to launch events may stabilize adapted priors, thereby limiting a full recovery even when the adaptor is removed. Additionally, recovery dynamics may vary across individuals. If individuals differ in the speed with which they return towards baseline, the recovery process may not be fully captured within the current paradigm.

A natural question is whether the recovery profile from adaptation would have looked different if observers had been deprived of *all* visual input between the adaptation and post-adaptation blocks. One way to achieve this would have been to ask observers to close their eyes during the break, which would rule out any possibility of visual input. This would have provided a more stringent control than our podcast break, during which no changing visual stimuli were presented but observers still had access to a neutral display. We chose not to adopt this stricter control because it seemed too burdensome for observers to sit in a dark room with closed eyes for an extended period. While we acknowledge that an eyes-closed condition would have been a cleaner methodological control, we consider it unlikely that it would alter the results. Given that we already observed an instananeous recovery following the podcast break, it seems improbable that a slightly more extreme form of blocking visual input would produce a qualitatively different recovery profile.

Visual adaptation proved to be an effective and robust tool to address how spatially specific adaptation to causality is, and how the perception of causality recovers from adaptation. We provide compelling evidence that the mechanisms for processing causal interactions are highly spatially localized, rapidly responding to the onset of adaptation and then quickly recovering in a time-based manner, eventually yielding nearly complete recovery that can feature slight differences to the pre-adaptation level. The combination of pronounced spatial specificity and instananeous recovery supports our idea that the interpretation of causal interactions originates at an early, perceptual stage of processing.

## Acknowledgments

This research was supported by a DFG research grant to S.O (OH 274/4-1) as well as funding from the Heisenberg Programme of the DFG to S.O. (OH 274/5-1). The authors declare no competing financial interests.

